# Reproducibility and model-selection stability in connectome-constrained circuit modeling

**DOI:** 10.64898/2026.04.18.717873

**Authors:** Christos Karaneen, Erik W. Schomburg, Dmitri Chklovskii

**Author notes:** **Corresponding author** Christos Karaneen.

## Abstract

Connectome-constrained neural network models aim to link anatomical connectivity with functional computation by training networks whose architectures reflect biological circuits. Because such models are increasingly used to infer neural mechanisms, it is important to assess their robustness to variations in training conditions and model selection criteria. Here we retrain ensembles of connectome-constrained models under nominally identical conditions and compare their correspondence to experimentally measured response properties in the *Drosophila* motion pathway. While task performance remains similar across models, the identification of biologically plausible circuit solutions is unstable across retraining runs. In particular, model clusters selected by lowest validation task error do not reliably correspond to experimentally observed neural tuning, and small variations in performance metrics can reorder cluster rankings. These results indicate that, in this framework, similar task performance does not reliably identify biologically plausible circuit solutions. Task error alone is therefore insufficient for mechanistic identification, and additional model-selection criteria are needed.

## 1 Introduction

Connectome-constrained neural network models aim to infer neural computation from anatomical connectivity. The framework introduced by Lappalainen et al. (2024) constructs neural networks whose architecture is strongly constrained by the *Drosophila* optic-lobe connectome, and optimizes the remaining parameters on a motion-detection task. The authors then compared the resulting responses of neurons within the model networks to stimuli known to elicit particular responses in physiological experiments, and found that top performing networks in the motion detection task tended to have unit responses similar to those observed in the corresponding biological neurons. Here we examine the robustness of such inferences to retraining and model-selection choices.

In our hands, the framework reproduced many qualitative outcomes reported in the original study, but the functional and mechanistic insights derived from “best-performing” models were not stable across nominally identical training runs. In particular, the different model clusters, some of which exhibited similar neural responses to biology while others did not, had overlapping distributions of validation task error, and the lowest-error models did not consistently match experimentally observed response tuning across retraining runs. This matters because, when multiple distinct circuit implementations achieve similar validation error, the objective function itself cannot arbitrate between them — and mechanistic interpretation becomes contingent on selection choices that are underdetermined by the training procedure.

## 2 Results

We focused on the motion pathway because it provided one of the clearest biological validation cases in the original study. T4 and T5 neurons in the *Drosophila* optic lobe are well-characterized direction-selective cells whose tuning to motion direction and stimulus polarity has been measured experimentally. These response properties serve as a concrete benchmark: a biologically plausible model should recover the experimentally observed tuning curves in these neurons.

In the original ensemble, clustering models by T4c activity similarity yielded a cluster whose members best matched experimentally measured T4c neuron tuning, and this same cluster also had the lowest mean validation task error (Fig. 1a). The original study highlighted this association to illustrate how biologically realistic tuning correlates with lower task error in this setting. More broadly, the 10 lowest-error models showed good correlations between the flash responses and moving-bar tuning curves of numerous cells in the motion pathway and their experimentally measured responses (Fig. 1b).

**Fig. 1.**
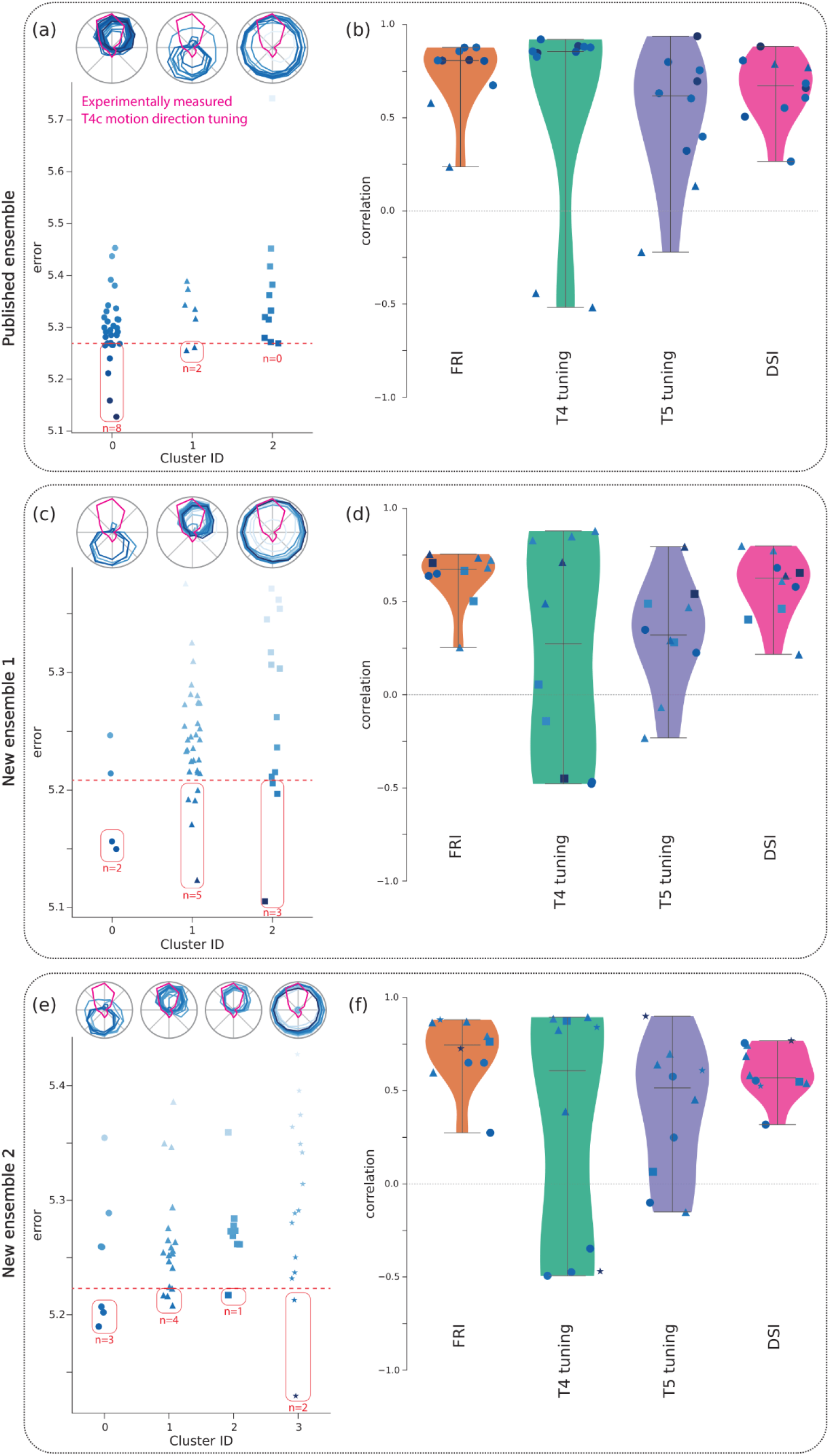
Two newly trained ensembles of connectome-constrained network models exhibit higher variability and an unreliable relationship between task-based model selection and biological plausibility. **a**, In the published ensemble, after clustering the network models by T4c activity similarity, the cluster with most of the best-performing networks exhibited accurate motion direction tuning in T4c neurons. **b**, Overall, the top 10 (20%) of models by task performance tended to have strong correlations with experimentally measured stimulus responses. **c**, A retrained network ensemble showed weaker relationships between task error and experimentally observed response tuning, with biologically matched model clusters being less easily selected with task performance-based criteria. **d**, Across cell types, correlations with experimentally observed response tuning were weaker and more variable. **e, f**, A second retrained network exhibited similarly degraded results.

When we trained two new ensembles of 50 networks, however, under nominally identical training conditions, the cluster with the lowest mean validation error did not reliably correspond to the best match with experimentally observed response features (Fig. 1c,e; Supplementary Figs. 2–3). Task errors varied only modestly across models — cluster mean errors differed by less than 0.08 across both new ensembles — so that differences in validation error were often too small to reliably distinguish between candidate circuit solutions.

Alternative selection rules could sometimes recover clusters with better biological agreement, but these rules were ad hoc, only marginally more successful, and not robust to small variations in performance metrics. For example, selecting the cluster containing the largest number of networks among the top 20% ranked by task error identified the cluster that most closely reflected experimental observations (Fig. 1a,c,e; Supplementary Figs. 1b–3b), yet this criterion was arrived at post hoc by inspecting which rule recovered the desired cluster, rather than from an independent theoretical motivation — and small changes in the task error could have changed which activity cluster would have been selected as the best performing one. Furthermore, in our retrainings, correlations between top-ranked models and experimentally characterized tuning were weaker and more variable (Fig. 1d,f). Together, these observations indicate that task performance alone is insufficient for reliable mechanistic identification when multiple solutions achieve similar error.

## 3 Discussion

More broadly, these results show that for connectome-constrained models, mechanistic conclusions depend not only on architecture and training objective, but also on how models are selected from among comparably performing solutions. This does not preclude the possibility that more detailed circuit models or alternative objective functions could improve robustness. For example, activity-based self-supervised objectives have shown promise in connectome-constrained models of other *Drosophila* circuits (Duan et al., 2025), and self-supervised learning on natural stimuli can recover direction-selective tuning and synaptic weight patterns consistent with connectomic data in the motion pathway, without supervised task objectives or error backpropagation (Qin et al., 2025). This nonetheless underscores the need to evaluate reproducibility and model-selection stability for any specific framework on which mechanistic claims are based.

In correspondence, the authors clarified that the paper’s primary accuracy claims rest on the median responses across the top 20% of models ranked by validation error, and that the clustering analysis is exploratory. However, it remains unclear how such median responses should be constructed and interpreted for mechanistic purposes, because they need not correspond to any single realizable network. Because mechanistic interpretation ultimately requires identifying specific, realizable circuit implementations, reliance on ensemble medians complicates biological inference.

Our aim is therefore not to argue that connectome-constrained modeling is uninformative, but to clarify what additional evidence is needed for robust mechanistic interpretation. On this basis, we believe two clarifications would substantially strengthen interpretation and reuse of the framework. First, principled and quantitative criteria for selecting biologically plausible clusters or models beyond validation task error would be valuable. Task error alone appears insufficient to uniquely identify biologically plausible circuit implementations when multiple solutions achieve similar performance, and additional diagnostics—such as sensitivity analyses, multi-task or activity-based constraints, out-of-distribution tests, or objective functions better aligned with circuit dynamics—may help avoid selection rules that are post hoc with respect to the biological ground truth.

Second, although the paper analyzes an ensemble of 50 models, more detailed characterization of variability across retraining runs, sensitivity of conclusions, and justification of ensemble size would improve interpretability. Full specification of random number generator seeds and elimination of nondeterministic training elements would enable exact reproducibility and systematic follow-up investigations. Finally, because the framework depends on a specified training objective and decoder, explicitly characterizing how mechanistic conclusions depend on these choices would strengthen generalizability.

We appreciate the impact this work has had in motivating connectome-constrained approaches to circuit modeling, and we hope these observations will help improve the framework’s robustness and interpretability for the broader community.

## 4 Methods

Retraining experiments were performed using the publicly available flyvis repository and training procedures described in Lappalainen et al. (2024), without modification to the training objective, hyperparameters, or decoder architecture. Two new ensembles of 50 networks each were trained with different random initialization seeds under otherwise identical conditions. Clustering of trained networks by T4c activity similarity and UMAP dimensionality reduction followed the same procedure as the original study. Biological correspondence was assessed by correlating model flash response indices (FRI), T4 and T5 motion direction tuning curves, and direction selectivity indices (DSI) with experimentally measured values, as in the original study.

## Code availability

Retraining experiments used the flyvis repository (https://github.com/TuragaLab/flyvis). Code used for the analyses in this work is available at https://github.com/eschombu/flyvis/tree/matters-arising.

## Contributions

C.K., E.S., and D.C. conceived the analysis. C.K. performed the retraining experiments. C.K. and E.S. analyzed results. All authors contributed to writing and revision of the manuscript.

## Acknowledgments

This work was supported by the Simons Foundation.

