## Supplementary Figures for "Reproducibility and model-selection stability in connectome-constrained circuit modeling"

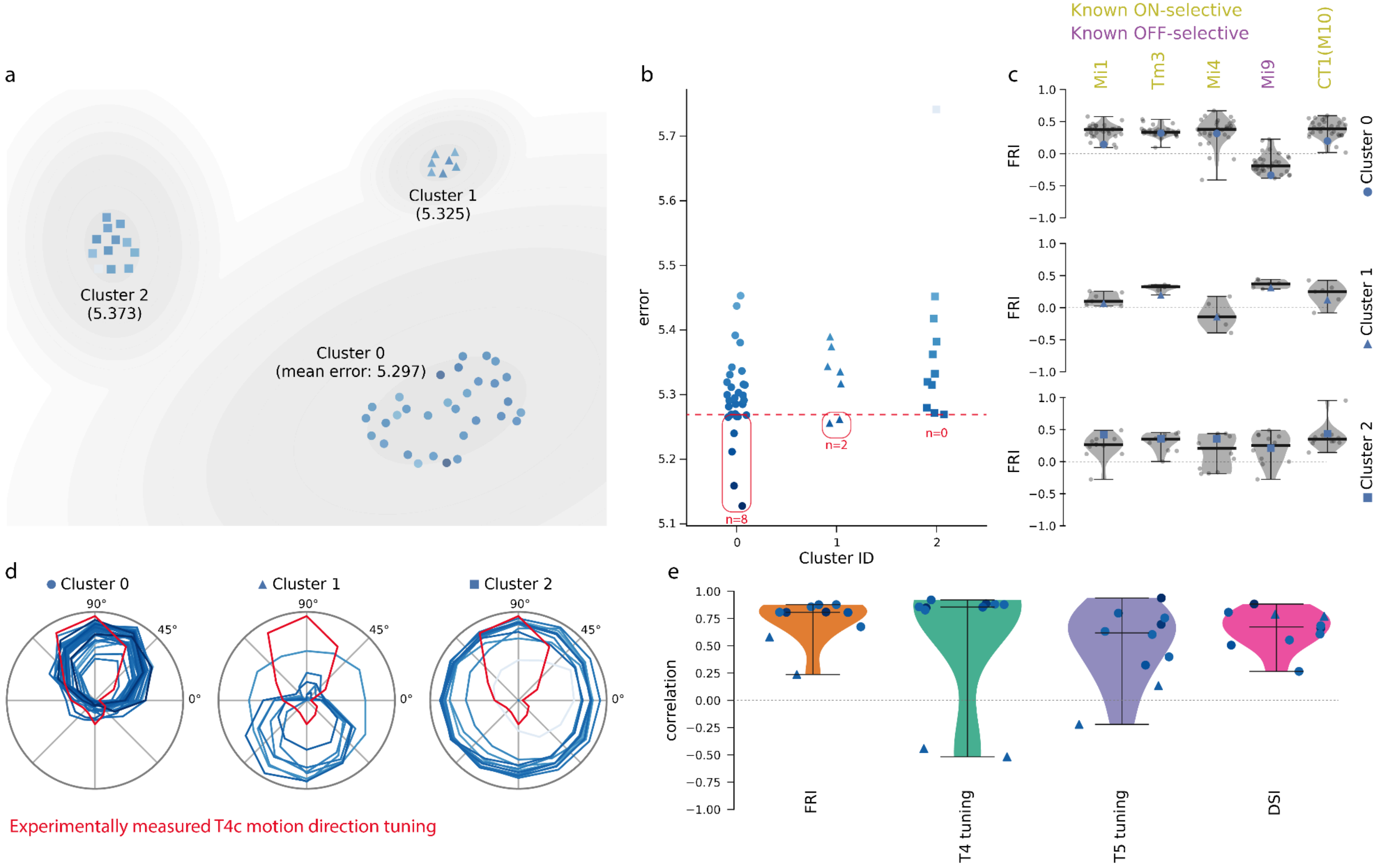


**Supplementary Fig. 1. ON/OFF flash responses and motion direction-selectivity of the best cluster of networks reflect experimental observations in the published results.** The results reported in Lappalainen et al. (2024) show that the best performing network clusters in the ensemble corresponded to experimentally observed responses to visual stimuli. **a,** UMAP projection of networks by T4c activity measures, showing three clusters, along with the mean validation task error for each cluster. **b,** The validation task errors for each network, segregated by cluster. Red dashed line and boxes highlight the 10 best models by task error. **c,** Flash response index (FRI) values for selected T4-input neurons. Cluster 0, which had the lowest mean task error, showed a good correspondence with known ON and OFF stimulus selectivity in Drosophila. **d,** The motion direction tuning curves for T4c neurons in each network, segregated by cluster. Cluster 0 again reflected experimentally observed motion direction tuning (Maisak et al., 2013). **e,** The correlations for the top 10 networks, ranked by validation task error, between: (i) model FRI values vs. known ON/OFF selectivity across cell types; (ii) model T4 and T5 edge motion direction tuning vs. experimentally measured tuning curves; and (iii) model direction selectivity index (DSI) vs. known ON/OFF edge motion selectivity across cell types.

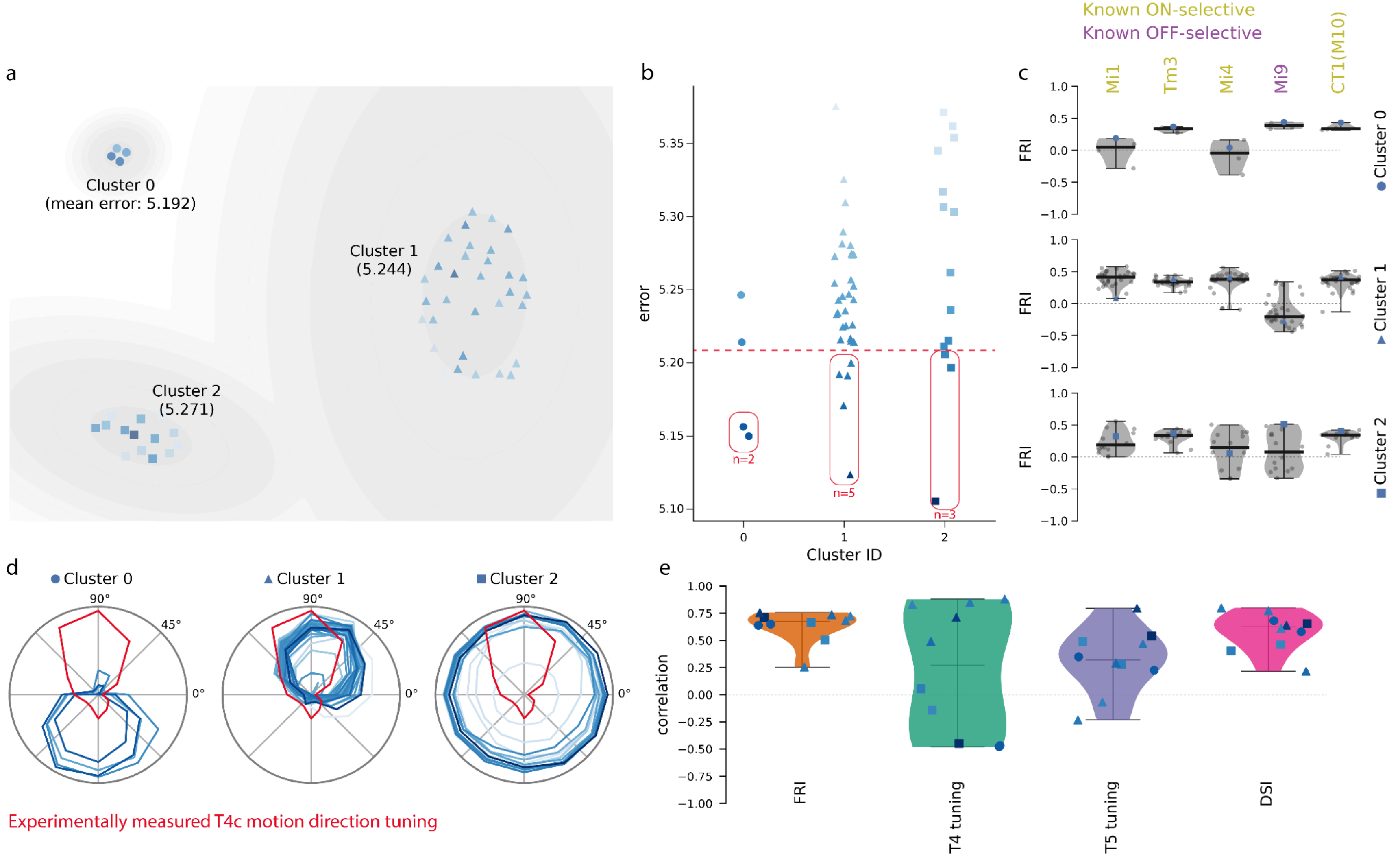


**Supplementary Fig. 2. The lowest-error models and model cluster do not reflect experimentally observed response tunings in a newly trained network ensemble. a,** In the newly trained network ensemble, a small cluster (4 networks) had the lowest mean validation error of the three clusters. **b,** The lowest mean-error cluster was cluster 0, and the best performing network was in cluster 2. Cluster 1, however, which matched experimentally observed responses, had many more members and more members within the 10 best-performing networks. **c, d,** The networks in cluster 0 exhibited a much weaker correspondence to experimentally measured neural responses, including FRIs and T4 ON-edge motion direction tuning. **e,** The correlations of neural responses were generally worse for the 10 best performing models in this ensemble.

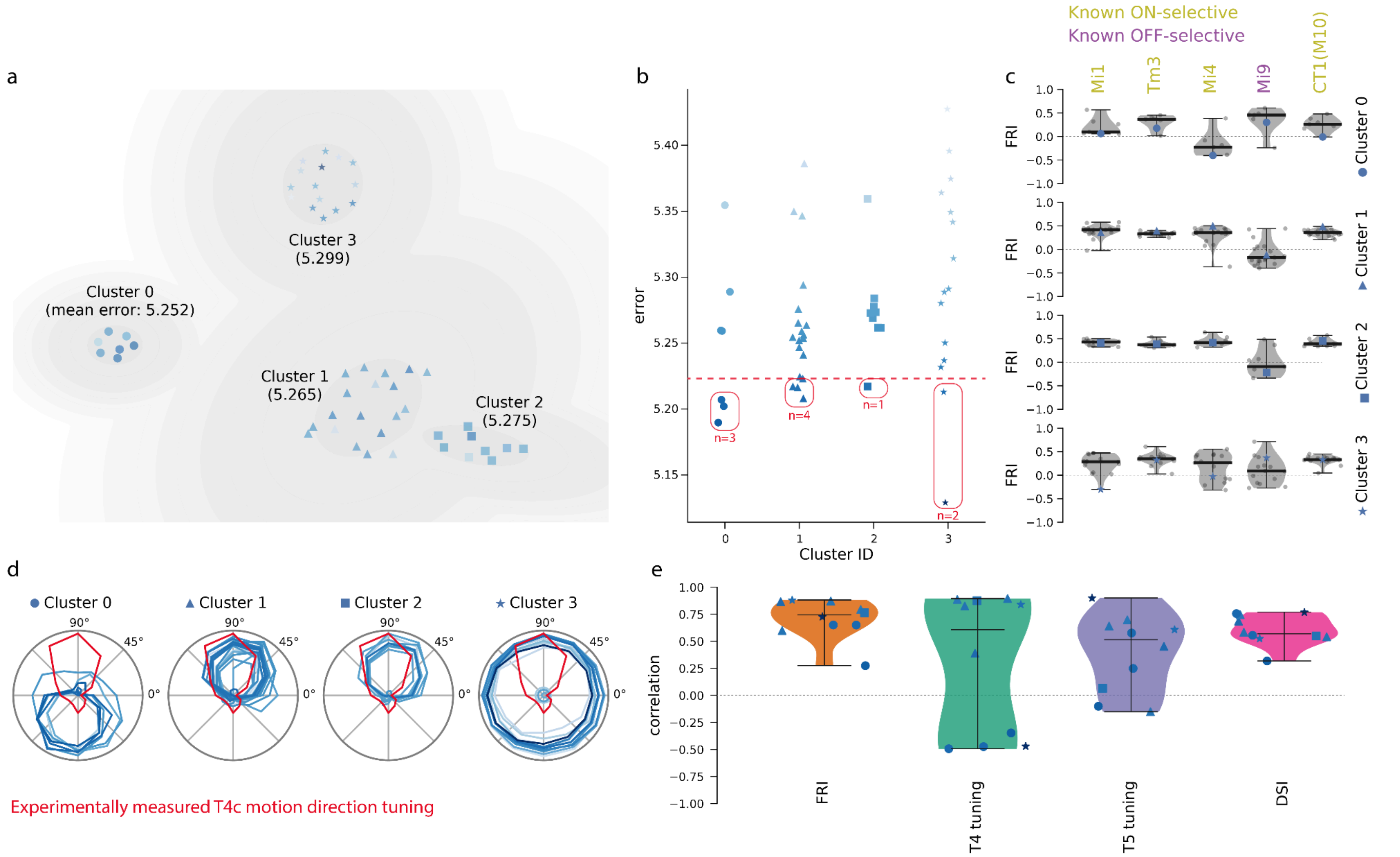


**Supplementary Fig. 3. Similarly worse results for a second newly trained network ensemble. a,** A possible fourth cluster, though with similar performance and neural activity / responses as its nearest neighbor cluster, emerged in a second newly trained ensemble. **b-d,** Again, the best performing cluster and model did not exhibit a good match to experimental observations of neural responses to ON/OFF stimuli. A cluster selection criterion of the highest count of networks in the best 10 / best 20% of models could again choose the cluster reflecting experimental observations, but this criterion is neither especially principled, nor robust to small variations in performance metrics.
